# Leafhopper Viral Pathogens: Ultrastructure of salivary gland infection of Taastrup-like virus in *Psammotettix alienus* Dahlbom; and a novel Rhabdovirus in *Homalodisca vitripennis* (Germar) Hemiptera: Cicadellidae

**DOI:** 10.1101/465468

**Authors:** Thorben Lundsgaard, Wayne B. Hunter, Scott Adkins

## Abstract

Viruses that are pathogenic to insect pests can be exploited as biological control agents. Viruses that are pathogenic to beneficial insects and other arthropods, as in honey bees, silk worms, and shrimp, cause millions of dollars of losses to those industries. Current advances in next generation sequencing technologies along with molecular and cellular biology have produced a wealth of information about insect viruses and their potential applications. Leafhoppers cause economic losses as vectors of plant pathogens which significantly reduce the worlds’ food crops. Each year more viruses are discovered primarily through the use of next generation sequencing of the leafhopper hosts. The diversity of viruses from leafhoppers demonstrates a wide range of taxonomic members that includes genomes of DNA or RNA from families like: Reoviridae, Iridoviridae, Dicistroviridae, Iflaviridae, and others yet to be classified. Discussed is a recent viral pathogen isolated from the leafhopper *Psammotettix alienus*, name Taastrup Virus. Taastrup virus (TV) is a novel virus with a RNA genome, a Filovirus-like morphology, being tentatively placed within the *Mononegavirales*. Adult *Psammotettix alienus* infected with TV, showed the highest concentration of virions in salivary glands, consisting of a principal gland (type I-VI-cells) and an accessory gland. Examination of thin sections revealed enveloped particles, about 1300 nm long and 62 nm in diameter, located singly or in paracrystalline arrays in canaliculi of type III- and IV-cells. In gland cells with TV particles in canaliculi, granular masses up to 15 μm in diameter were present in the cytoplasm. These masses are believed to be viroplasms, the sites for viral replication. TV particles were observed at the connection between a canaliculus and the salivary duct system. A TV-like virus with strongly similar morphology was discovered in the ornamental plant, *Liriope*, near Fort Pierce, Florida, USA. When the virus was inoculated to a leafhopper cell culture, HvWH, made from the glassy-winged sharpshooter, *Homalodisca vitripennis* (Germar), the cells rapidly degraded with 100% mortality in 48 hours. These two instances are the only reported cases of this newly discovered viral pathogen of leafhoppers.

## 1. Introduction

Insects are commonly infected with multiple viruses and traditional methods of detection and isolation, such as observation of sick or dead insects in the field can miss many pathogens as diseased insects fall to the ground and are quickly scavenged by ants or other predators. The use of insect cell cultures to ‘capture’ virus by inoculation of crude preparations made from insects collected from the field, can propagate virus if the cell lines are permissive to that specific virus (Hunter et al, 2001) however, these methods only capture a small fraction of the potential pool of viruses that are infecting insects and other arthropods. Mining of the genetic sequences now available, for new insect viruses, has proven to be a rich resource full of undiscovered, insect pathogens (Katsar et al, 2007; Valles et al, 2004, 2008; Liu, et al, 2011). Even so, serendipity still provides many new discoveries for the prepared mind. Recently a Taastrup-like leafhopper-infecting virus was discovered in the ornamental flowering plant, Genus: *Liriope*, Fort Pierce, Florida, USA (Fig. 1). The plant had a mixed infection with a Tospovirus that produced the traditional ring-like symptoms on the leaves (Fig. 2). During these examinations, a Taastrup-like virus was identified. Current research efforts are focused on identifying potentially new leafhopper pathogens which may have a function as biological control agents.

**Figure 1.**
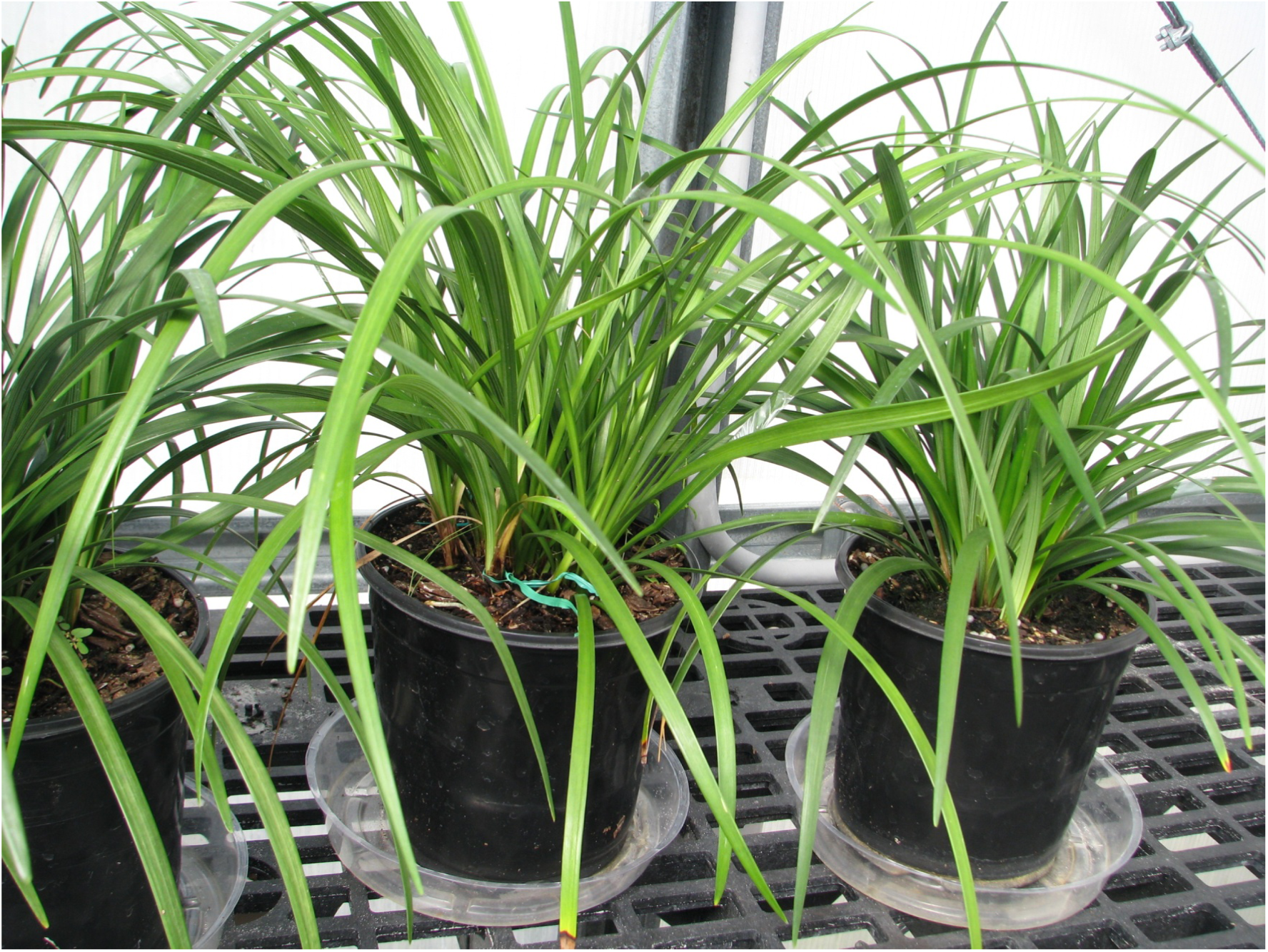
Healthy non-infected *Liriope* is a plant belonging to the lily family and is one of the most popular ground covers in Florida. It is also called ‘Lilyturf or ‘monkey grass’ and is a native plant of Eastern Asia.

**Figure 2.**
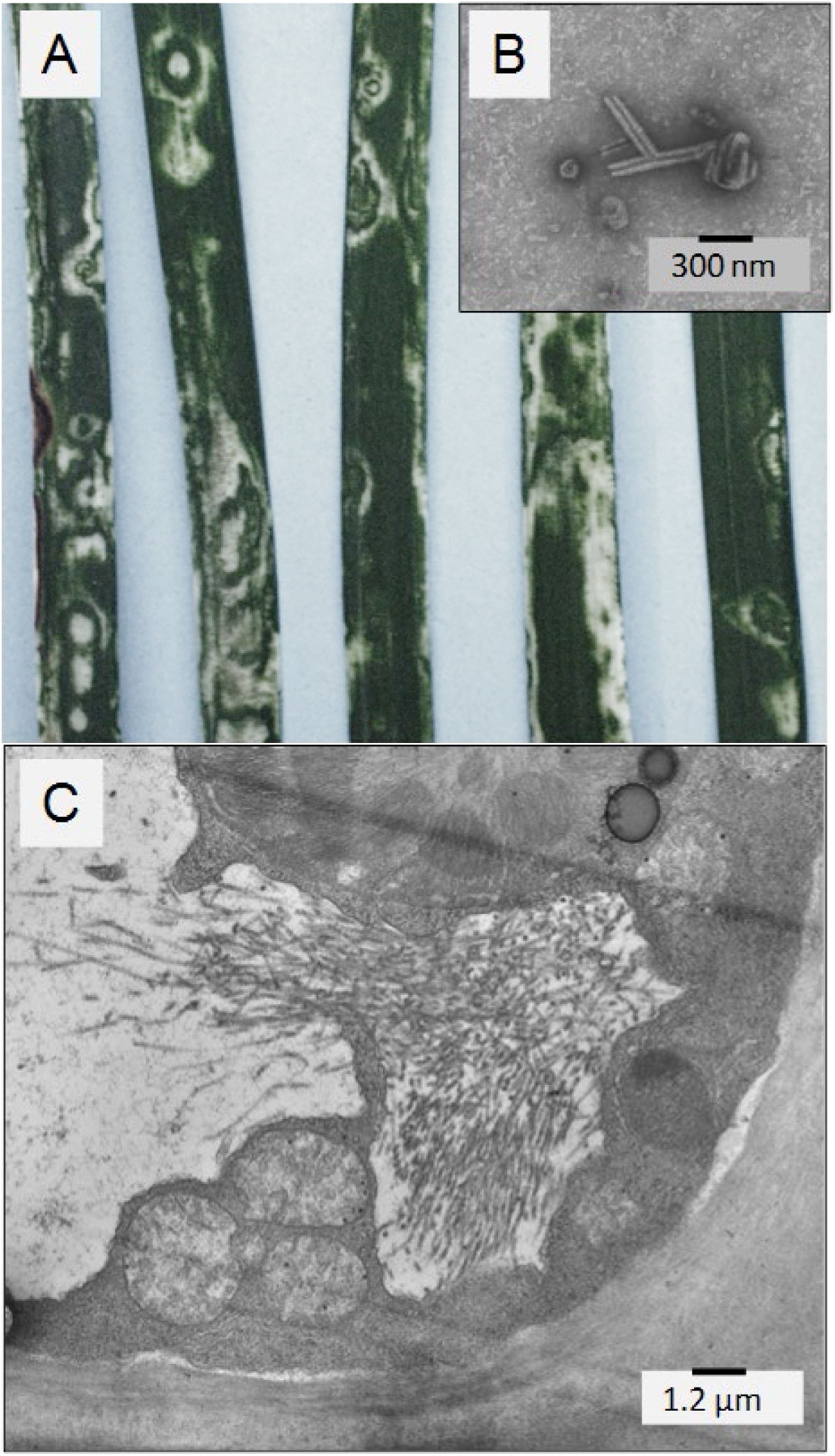
Flowering plant, genus, *Liriope*, with ring-like symptoms typical of a tospovirus infection-(A). Electron microscopy examination of plant tissues however, showed a mix infection with one virus having distinct morphology similar to Taastrup virus (B- Taastrup-like virus Florida strain); Plant tissues showed typical virus infection associated with the plant-infecting Rhabdoviruses (C).

Rhabdoviruses (Family: Rhabdoviridae) in the Order *Mononegavirales*, are enveloped viruses with a nonsegmented negative-strand RNA genome. Currently this Order includes four families—Rhabdoviridae, Bornaviridae, Filoviridae, and Paramyxoviridae. Rhabdoviruses have a diverse host range including humans, vertebrate animals, and invertebrates in the aquaculture and agricultural industries. The International Committee on Taxonomy of Viruses, ICTV, recognizes nine genera in the family Rhabdoviridae (Dietzgen et al. 2011). These include viruses considered to be plant viruses transmitted by arthropod vectors of the genera *Nucleorhabdovirus* (10 species) and *Cytorhabdovirus* (9 species); with members in the genera *Ephemerovirus (5 species), Lyssavirus (12 species), Novirhabdovirus (4 species), Perhabdovirus (3 species), Sigmavirus (7 species)*, Tibrovirus *(2 species), Vesiculovirus (10 species), and one unassigned genus (6 species)*, which have been isolated from various vertebrate and invertebrate hosts (www.ictvonline.org/virusTaxonomy.asp). Although more than 150 rhabdoviruses have been identified from a wide range of hosts worldwide, most still remain to be assigned to a genus within a family due to the high amount of genetic diversity among them (Kuwata et al, 2011).

Filovirus-like particles of Taastrup virus (TV) have been described from a population originating from France of apparently healthy leafhoppers belonging to the species *Psammotettix alienus* Dahlbom (Hemiptera: Cicadellidae)(Lundsgaard, 1997). *Psammotettix* leafhopper species are extensively listed in the literature as feeding on grasses and occur worldwide. The type specie of the Genus, *Psammotettix maritimus* (Penis), feeds on *Convolvulus* (Haupt, 1929; DeLong, 1973). Examination of plant and leafhopper tissues showed flexuous particles, 55-70 nm in diameter and 600 or 1100 nm long, consist of an inner coiled nucleocapsid about 30 nm in diameter and surrounded by a membrane with pronounced surface projections inserted (Lundsgaard, 1997), thus resembling members of Filoviridae morphologically. The RNA genome has partly been sequenced, which demonstrated both, that TV belongs to the virus Order *Mononegavirales*, but which has not been officially classified into any of the established families of this virus order (Bock *et al*., 2004). However, TV isolated from the leafhopper *P. alienus*, in Denmark, AY423355, was inferred to belong to the family Rhabdoviridae, in the genus *Cytorhabdovirus* based on analyses of the L polymerase gene (Bourhy et al., 2005). The hosts for *P. alienus* are grasses and cereals (Raatikainen and Vasarainen, 1976). The insects salivate during probing, penetration, and plant fluid ingestion (Backus, 1985), thus it is likely that the salivary glands play a key role in the epidemiology of TV. Thus, a more in-depth study was conducted on the ultrastructure of organs from infected leafhoppers with results presented here.

## 2. Experimental Section

The following experiments were performed in order to determine the distribution of virus in leafhoppers taken from a TV-infected population. Two fractions were made from ten randomly chosen insects. One fraction consisted of heads (including salivary glands) and the other fraction of thorax, abdomen, wings and legs. Each of the fractions was extracted in 1 ml distilled water. After one cycle of differential centrifugation (5 min at 1800 *x* g and 20 min at 13,000 *x* g), the sediments were dissolved in 20 μl (head fraction) or 100 μl (body fraction) of negative staining solution (0.5% ammonium molybdate, pH 7). Preparations were made on Formvar^®^ mounted nickel grids (400-mesh) and the number of TV particles in ten grid squares was counted in a transmission electron microscope (JEOL JEM-100SX). The mean number of particles for the head fraction was 9.1 [standard deviation (SD) = 4] and zero particles for the body fraction. In the next experiment, the heads from ten insects were severed and two fractions prepared, namely: a salivary gland fraction (SG) and a fraction representing the remaining part of the head (H). After differential centrifugation, both sediments were dissolved in 20 μL staining solution. The mean number of particles per grid square was 34.3 (s = 5) for SG and 0.3 (s = 0.7) for H, showing the salivary glands to be the most important site for accumulation of TV particles.

Ten adults taken in random from the TV infected population were dissected in Ringer’s solution (0.7% NaCl, 0.035% KCl, 0.0026% CaCl_2_). Each pair of salivary glands was divided in a left and a right part with the purpose of checking one part for presence of particles by negative staining electron microscopy (extraction in 10 μl 0.5% ammonium molybdate) and the other corresponding part by ultrastructural analysis. The salivary glands were fixed, dehydrated, and embedded essentially according to Berryman and Rodewald (1990). In brief, the specimens were transferred to fixative (3% formaldehyde, 0.3% glutaraldehyde, 100 mM phosphate buffer, pH 7.0) and maintained in fixative for at least 2 h. After washing (3.5% sucrose, 0.5 mM CaC1_2_, 100 mM phosphate buffer, pH 7.4), the aldehyde groups were quenched with 50 mM NH_4_C1 dissolved in washing solution) for 1 h. The specimens were then washed (3.5% sucrose, 100 mM maleate buffer, pH 6.5) and post-fixed with 2% uranyl acetate in the same maleate buffer for 2 h. After dehydration in acetone (50%, 70%, 90%), the specimens were infiltrated in LR Gold (The London Resin Co., England) containing 0.5% benzoin methyl ether (3 changes) and polymerized under near ultraviolet light (Philips TW6). Fixation and dehydration (including 50% acetone) was done at 5-10 °C and dehydration (from 70% acetone), embedding, and polymerization performed at - 20°C. Thin sections (60-70 nm) were collected on 200-mesh nickel grids, stained with saturated uranyl acetate for 5 min and lead citrate (1 mM, pH 12) for 1 min, and examined with a JEOL 100SX transmission electron microscope operated at 60 kV. As controls, specimens were prepared in the same way from a healthy population of *P. alienus*.

Field collected *Liriope* which had symptoms of a tospovirus-like infection (Figs. 1, 2) were prepared for examination by TEM as mentioned above. Upon discovery of a Taastrup-like virus infection a crude sap inoculum was prepared. Solutions were prepared from *Liriope* plants with and without symptoms and the resulting inoculum dispensed onto leafhopper cell cultures produced from the glassy-winged sharpshooter, *Homalodisca vitripennis*, HvWH, Hunter, USDA. The crude plant sap preparation used 2 g of leaf tissues homogenized in Histidine, monohydrate buffer, pH 6.5, which was the same buffer used for insect cell cultures (Hunter and Polston 2001; Hunter et al, 2001; Marutani et al, 2009). The solution was then centrifuged in 15 mL tubes at 600 xg for 4 min to pellet the plant debris. The supernatant was collected and filtered through a 0.45 μm filtration unit, and then filtered again through a 0.22 μm filtration unit. The filtered sap inoculum was then added to adherent leafhopper cell cultures at 1 mL per 25 cm^2^ flask (Corning®, Inc), and let stand for 10 minutes. After this time 4 mL of fresh insect medium was added (Table 1). Leafhopper cell cultures were observed daily for cell pathogenicity.

**Table 1.** Hunter’s Leafhopper Cell Culture Medium, H2G, components. [**1A, B**] *Homalodisca vitripennis* (Germar) (Hemiptera: Cicadellidae) cells in culture started from embryonic tissue from eggs. Adult glassy-winged sharpshooter leafhopper.

|  |  |
| --- | --- |
| Grace's Insect medium (supplemented, 1X) | 210 mL |
| 0.06M L-histidine monohydrate buffer solution pH=6.5 | 290 mL |
| Medium 199 (10X) | 10 mL |
| Medium 1066 (1X) | 17 mL |
| Hank's Balanced Salts (1X) | 33 mL |
| L-Glutamine (100X) | 1.5 mL |
| MEM, amino acid mix (50X) | 1.5 mL |
| Pen-Strep (w/ Glutamine) | 2.5 mL/500 mL |
| Nystatin | 1.0 mL/500 mL |
| Gentamycin | 1.5 mL/500 mL |
| Dextrose | 1.8 g |
| Fetal Bovine Serum | 10% of final volume |

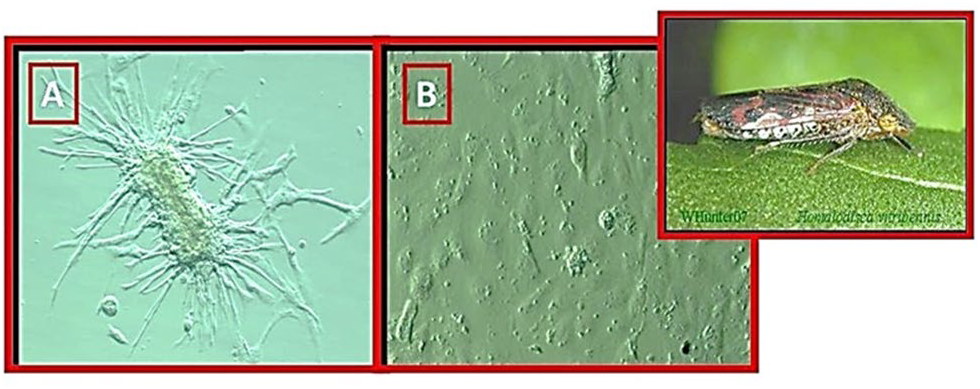

Leafhopper cell line maintenance: HvWH, were cultivated in H2G Leafhopper medium, modified from WH2 honey bee media (Hunter 2010) (Table 1). Culture methods in (Kamita et al, 2005; Biesbock et al, 2014). Prior to adding the fetal bovine serum, the medium was passed through sterilizing filtration units (500 mL, 0.22 μm filter, Corning), then fetal bovine serum was added to make a 10% concentration. Aliquots of 5 mL of medium were placed on the counter for three days at room temperature to test for bacterial contamination. Fungin™ a new soluble formulation of Pimaricin (InvioGen, Cat. No. ant-fn-2, 200mg) was added to the culture medium to inhibit mold growth. Cultures were maintained in Corning 25 cm^2^ tissue culture flasks treated with CellBIND™ to promote cell attachment (Corning®, Lowell, MA) (Hunter 2010). Culture flasks were kept in an incubator at 24°C and examined using an inverted microscope (Olympus IX70) at 10x, 20x, 60X magnifications. Complete medium change was done every 10 days without disturbing the culture surface and cultures were passed when approximately 80% confluent. Cell passage used 0.25% Trypsin EDTA solution (Invitrogen™, Carlsbad, CA) to dissociate cells by exposure for 2 to 5 min to achieve complete dissociation. Adherent cells had medium gently pipetted across the culture surface to detach cells. Once cells were dissociated, they were collected into 15 mL centrifuge tubes and centrifuged for three min at a force of 400xg, in a clinical centrifuge, (IEC, Centra CL2, Thermo Electron Corp., Milford, MA 01757). The supernatant was drawn off and the cell pellet gently resuspended in 4 mL of fresh medium. Cells were seeded at 2 mL per 25 cm^2^ flask, being split at a 1:2 ratio. Fresh medium was added to make a final volume of 5 mL per flask. Newly passed cultures were left untouched for 48 hours to let cells settle and attach to the flask substrate.

## 3. Results and Discussion

Sogowa (1965) has described the morphology of leafhopper salivary glands in detail, but *P. alienus* was not included and has not been described elsewhere. The morphology of these glands was therefore studied and described here. The naming of the glandular cells made by Sogowa (1965) is followed. The salivary glands of P. *alienus* are lying in the head and prothorax and consist of two pairs of glands, each made up of a complex principal gland (Fig. 3, I-VI) and an U-formed accessory gland (Fig. 3, AG). The lettering III, IV, and V all represents single cells, but the areas designated I, II, and VI in Fig. 3, represents one to more cells each (the exact number not determined). The accessory salivary gland (AG) consists of many cells. According to the ultrastructure of the canaliculi in type III-cells (see later), these cells are subdivided in four type IIIa-cells and two type IIIb-cells. The orientation of the principal salivary gland within the insect showed one IIIb-cell pointing downward and the other IIIb-cell against the front. The I- and II-cells (anterior lobe) form the proximal part and the IV- through VI-cells (posterior lobe) the distal part of the gland. The overall length of a typical principal salivary gland (from the tip of the lower IIIb-cell to the tip of the upper IIIa-cell) is about 550 μm.

**Figure 3.**
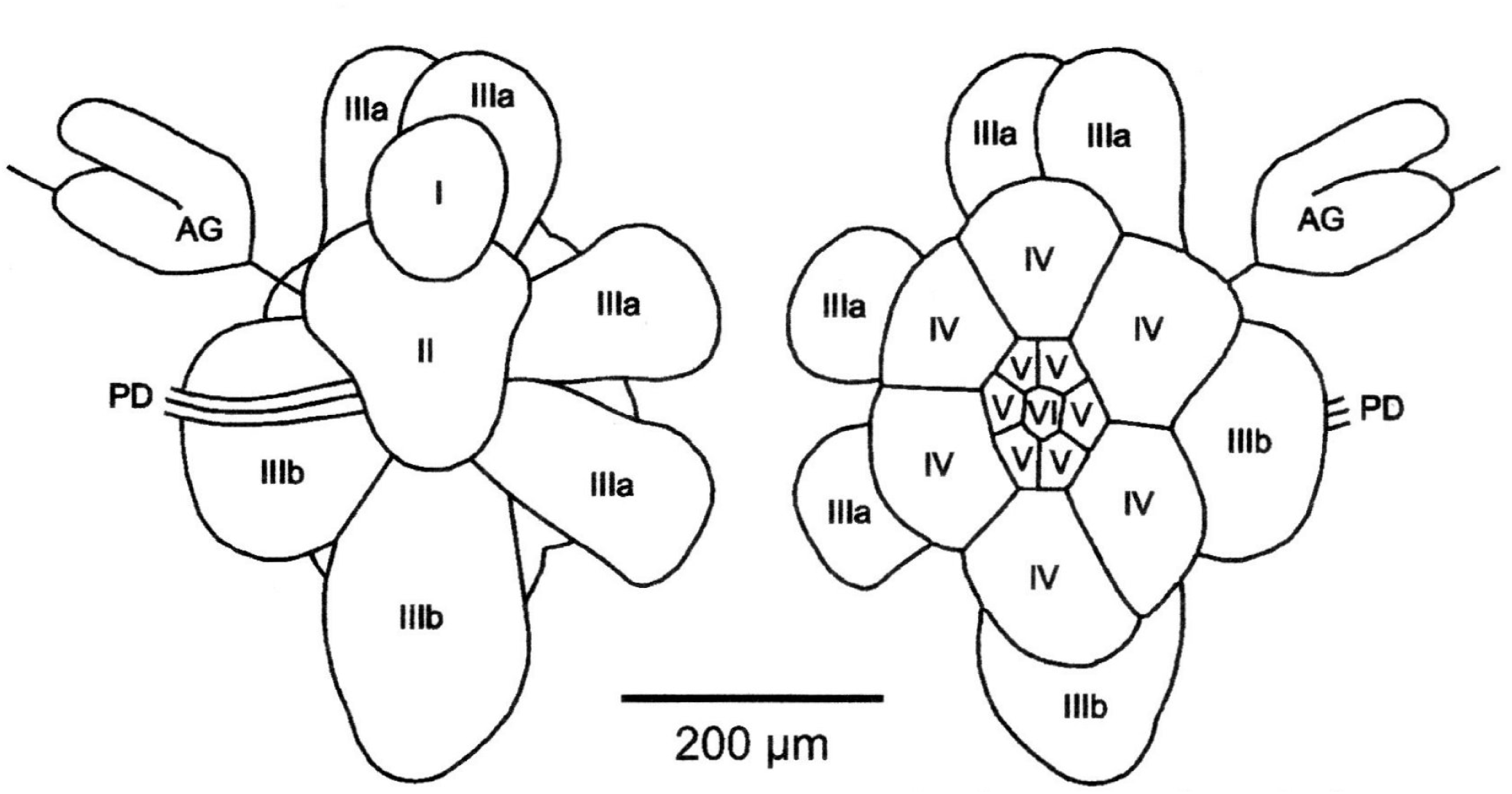
Drawing of salivary glands from *Psammotettix alienus* seen from the insect axis (left) or from outside (right). The salivary glands consist of a principal gland (cell type I to VI) and an accessory gland (AG), both connected to the principal duct (PD). According to ultrastructure of canaliculi, the type III-cells are differentiated in four type IIIa-cells (pointing upwards and backwards) and two type IIIb-cells (pointing forwards and downwards).

Negative staining electron microscopy revealed TV particles in glands from two of 10 examined insects. By ultrastructural analysis, unusual structures, as described below, were detected in glands from the same two individuals. Rod-shaped particles were observed either singly or in paracrystalline arrays in canaliculi of type III- and IV-cells (Fig. 4 -5). The type III-cells are easily distinguished from other salivary gland cells by not having secretory vesicles in the cytoplasm. However, two types of III-cells could clearly be distinguished in *P. alienus* according to the structure of canaliculi. In the one type, here designated IIIa, the content of the canaliculi is granular (Fig. 3 and 4A), whereas the other, designated IIIb, has canaliculi filled with many small membrane-bound spherules containing electron-opaque granules (Fig. 3 and 4B). The IV-cells could easily be identified from all other salivary gland cells by a visible line pattern in their secretory vesicles (Fig. 4C).

**Figure 4.**
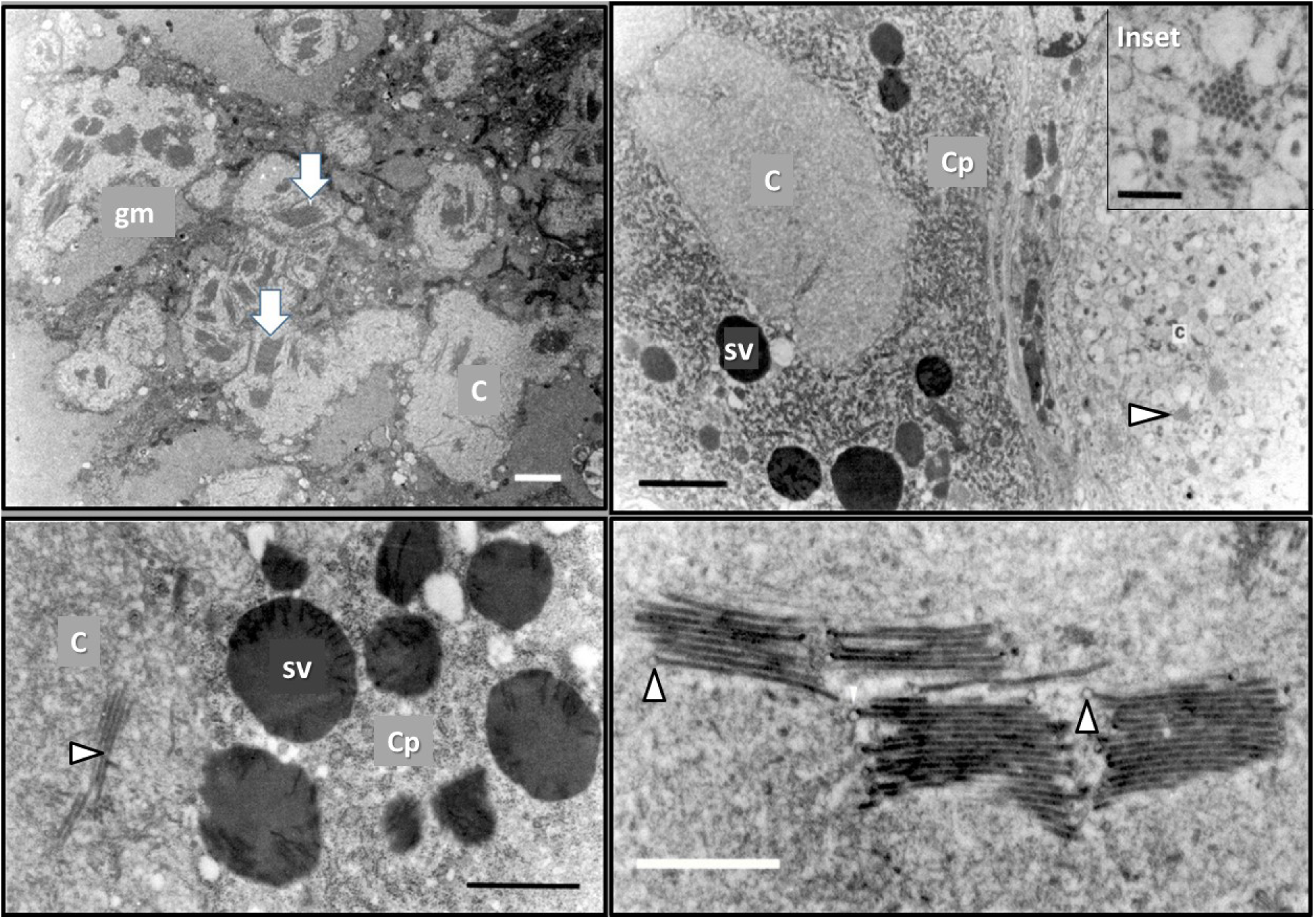
Section through a type IIIa-cell from *P. alienus* infected with Taastrup virus. Aggregates of virus particles (white arrowheads) are located in canaliculi (**c**) close to granular masses (**gm**) in the cytoplasm. Bar = 2 μm. [**4A**]; Section through a type IIIb- (right) and a type IV-cell (left) from *P. alienus* infected with Taastrup virus. Aggregates of virus particles (black arrowhead) are located in a canaliculus of the type IIIb-cell (inset). Note the vesicles with electron opaque granules in the canaliculus, characteristic of type IIIb-cells. The neighboring cell (type IV) has cytoplasm (**cp**) with rough endoplasmic reticulum together with electron opaque secretory vesicles (sv). No virus particles were observed in any of the canaliculi (c) of this cell. Bar = 2 μm. Inset bar = 500 nm. [**4B**]; Section through a type IV-cell from another infected leafhopper other than the one examined in Fig. 4. Virus particles (white arrowhead) are located in canaliculi (**C**) of this cell. Note the dark lines in the secretory vesicles (**sv**), which are characteristic for the type IV-cells. **Cp** =cytoplasm. Bar = 1 μm. [**4C**]; Longitudinal section through aggregates of virus particles located in a canaliculus of a type IIIa-cell from *P. alienus* infected with Taastrup virus. Small blebs (white arrowheads) are seen at one end of several particles. Bar = 1 μm. [**4D**].

The particles in the canaliculi were often seen with a membrane-like bleb at the one end (Fig. 4D). These blebs are probably remnants of viral envelopes remaining after a presumed budding process. For particles for which both ends could be seen, the mean length - not including the bleb - was calculated to be 1294 nm (s = 22). In cross section, an electron-opaque inner hollow core (about 31 nm in diameter) is seen surrounded by an outer faint electron opaque layer, separated by a translucent layer (Fig. 4D, 5A). The diameter of particles was calculated to be 62 nm from particles laying in paracrystalline arrays. The morphology and dimensions for the particles observed here in canaliculi are in agreement with the negative stained particles presented previously (Lundsgaard, 1997). This taken together with the correlation found between presence of particles in extracts and presence of particles in canaliculi among the 10 examined leafhoppers. The particles observed in canaliculi are identical to those described in the former paper. So, the inner, hollow core is probably the helical nucleocapsid of TV and the outermost faint layer the G protein spikes. In cross sections through particles of Rhabdovirus and Filovirus (Murphy and Harrison, 1979; Geisbert and Jahrling, 1995), an electron opaque layer, thought to be the virus membrane, is present between the layer of spikes and the inner nucleocapsid core. The analogous position of this electron opaque layer is translucent in the present TV particles. This discrepancy can be explained by use of osmium tetroxide for post-fixation (known to preserve and stain membranes) in the studies on rhabdo- and filoviruses, whereas osmium tetroxide was omitted in the present investigation. Osmium tetroxide was omitted in order to preserve the antigenic activity for future research on localization of viral antigens in these specimens. Aggregates of particles, 75-80 nm in diameter, were described from head cells of *P. alienus* (Lundsgaard, 1997). These particles were seen located in the cytoplasm (not in canaliculi as presented here) and their central, electron opaque core was not seen hollow (as are the present TV particles), suggesting that the particles described previously were not TV particles.

**Figure 5.**
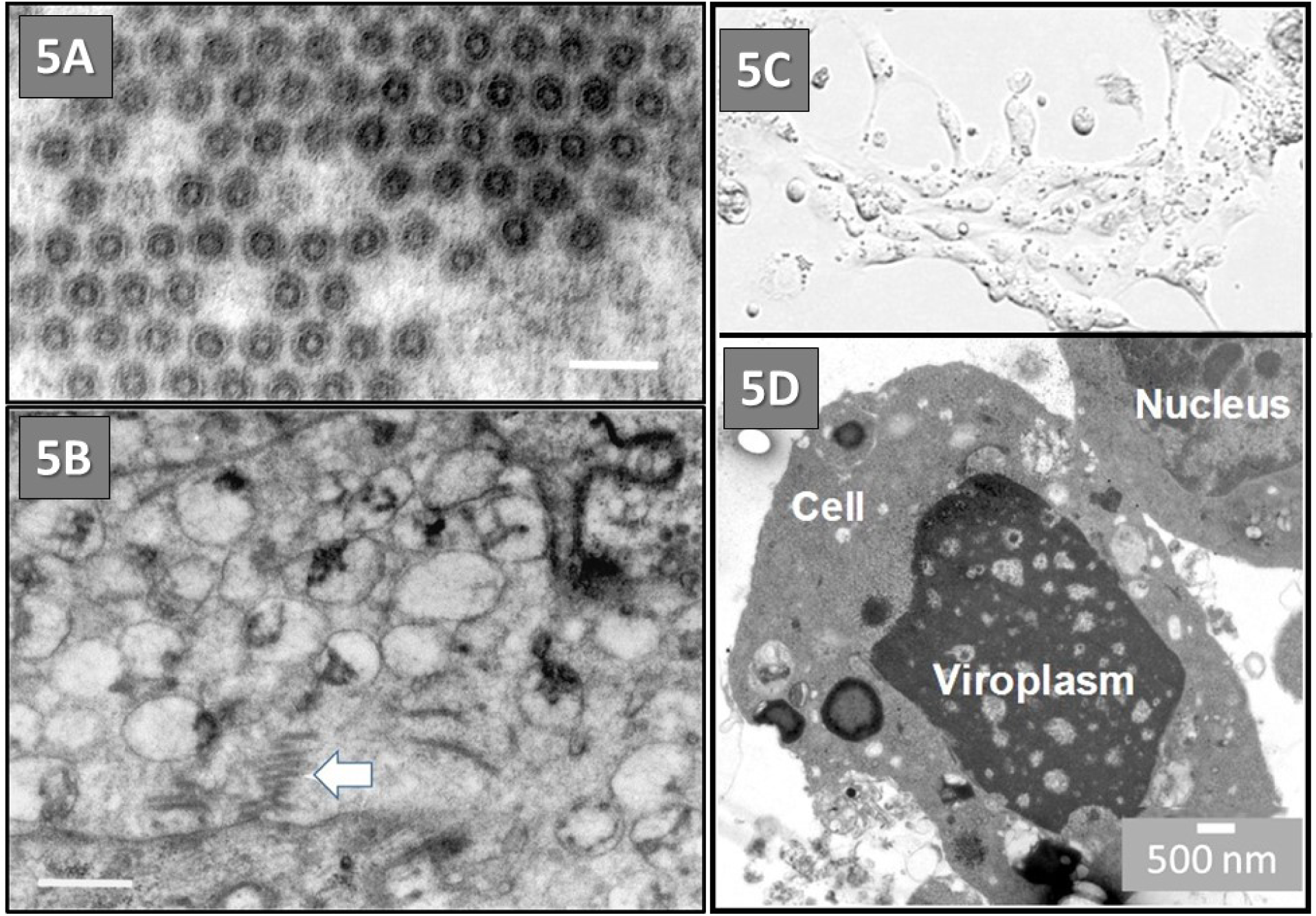
Transverse section through virus particles located in a canaliculus of a type IIIb-cell from *P. alienus* infected with Taastrup virus. An electron opaque tube with a hollow centre is seen surrounded by a translucent layer and outermost a faint electron opaque layer. TEM micrograph, Bar = 100 nm. [**5A**]; Close-up where a canaliculus opens into the salivary duct system. Oblique sections through some virus particles are seen (white arrowhead). Bar = 500 nm. [**5B**]; Light microscope image of Taastrup-like virus infected leafhopper cells *(H. vitripennis)* (**5C**) and transmission electron microscopy image (**5D**) of cell culture from glassy-winged sharpshooter, *Homalodisca vitripennis* (Germar), HvWH infected with a Taastrup-like virus (**5C**, 30x magnification) (Hunter and Adkins 2009). TEM of leafhopper cell with viroplasms-like structures in infected cells (**5D**, image provided by Diann Achor, University of Florida, Electron Microscopy Core, Lake Alfred, FL).

In the cytoplasm of type III- and IV-cells, possessing canaliculi with TV particles, fine granular masses up to 15 μm in diameter were observed (Fig. 4A). These masses do not contain ribosomes or other cellular organelles and their presence was positive correlated with presence of TV particles in nearby canaliculi. Granular masses (viroplasms), believed to be viral replication sites, have been observed in cells infected with Rhabdovirus (Murphy and Harrison, 1979; Conti and Plumb, 1977) and Filovirus (Geisbert and Jahrling, 1995). The granular masses present in the cytoplasm of salivary glands with TV particles are thus suggested to be viroplasms, the sites for TV replication. Because a nuclear localization signal has been identified in the G-protein gene of TV (Bock *et al*., 2004), nuclei close to viroplasms were carefully examined for abnormalities as compared with uninfected controls. Neither budding through nuclear membranes nor abnormality of nucleoplasm were observed during the present investigation.

In several cases, two glandular cells in contact with each other could contained both, viroplasms in the cytoplasm and TV particles in canaliculi, while the other neighboring cell was uninfected (Fig. 4B). These uninfected cells were of a type, which were infected elsewhere. TV particles were never seen between such cells or in the space between the plasma membrane and the basal lamina, so, the delivery of mature TV particles appeared only to take place by budding of nucleocapsids - synthesized in viroplasms - into canaliculi destined for saliva excretion. TV-particles or viroplasms were neither observed in type I-, II-, V-, and VI-cells of principal salivary glands, nor in accessory salivary glands from infected leafhoppers. TV-particles or viroplasms were not seen in salivary glands from leafhoppers originating from a TV-free population of *P. alienus*.

This investigation shows that salivary glands are the primary site of TV synthesis in adult *P*. *alienus*. One possible mode of TV acquisition relies on virions excreted with saliva into the plant tissue, which are passively delivered into the plant during feeding. Then TV is being ingested by uninfected nymphs and/or adults which become infected. Former investigations (Lundsgaard, 1997) have shown that host plants used for rearing TV infected leafhoppers, do not become infected, and so, the plants probably function as passive host acquisition for leafhopper vectors which then are able to transmit TV post replication latent period.

A Rhabdovirus, with strong similarity to the Taastrup virus was discovered in an ornamental plant, *Lirope spp*., grown in Florida, USA (Hunter and Adkins 2009; Hunter et al, 2009). A crude sap extract was prepared and screened on leafhopper cell cultures, *Homalodisca vitripennis* (Germar) glassy-winged sharpshooter (Hemiptera: Cicadellidae), for pathogenicity (Fig. 5C,D). The plant, *Liriope*, belongs to the lily family and is one of the most popular ground covers in Florida (Fig. 1). *Liriope* is a perennial, herbaceous, mat-forming plant that grows to about 30 cm tall. Leafhopper cell cultures inoculated with the virus-laden sap from infected *Liriope* plants resulted in massive cell death, 100% mortality in 48 hrs. Control cells inoculated with non-symptomatic plant sap inoculum showed no significant changes in cell morphology, adherence, or survival. Examination using TEM showed viroplasms-like structures in inoculated leafhopper cells treated with virus-infected *Liriope* sap, at 30 hrs post treatment (Fig. 5D). Attempts to sap inoculate non-infected *Liriope*, from the same field plots failed to produce symptoms. However, in the original *Liriope* plant with symptoms, the TV-like virions were observed at significant numbers in plant tissues to suggest replication (Fig. 2).

## 4. Conclusions

As genomic techniques continue to improve more viruses are rapidly being discovered (Babayan et al, 2018; Hunter et al, 2006; Hunnicutt et al, 2008; Liu, et al, 2011; Nouri et al, 2018; Rosario et al, 2018). The taxonomic group which includes Taastrup viruses and other related virus (Taastrup virus—Viruses; ssRNA viruses; ssRNA negative-strand viruses; Mononegavirales; Rhabdoviridae; Cytorhabdovirus; unclassified Cytorhabdovirus) continues to have new members discovered across a diverse range of insect Orders, from Hemiptera to Lepidoptera (Ma, et al, 2014). Insect infecting viruses are still examined for use as biological control agents (Lacey et al, 2015; Szewczyk et al, 2006), but applications of viruses have moved beyond just being pathogens. The advances in molecular biotechnology, low cost sequencing, bioinformatics software, and virus discovery pipelines to analyze large genetic datasets (Babayan et al, 2018; Bigot et al, 2018) have made virus discovery easier (Greninger 2018). Along with the rapid increase of virus discovery, has been the expansion of novel molecular biology applications for viruses, used in whole, or in part (promoters/peptides) across agriculture, human medicine, and material sciences (Hunter et al, 2019; Kolliopoulou et al, 2017; Wen and Steinmetz 2016). Virus can be used to delivery ‘genetic’ payloads, to produce RNA suppression of specific genes such as toxins in plants, or critical genes in an insect pest (Andrade and Hunter 2016, 2017; Kolliopoulou et al, 2017; Zotti and Smagghe 2015; Zotti et al, 2018). Since sap inoculation into uninfected plants was not successful, it may be that the virus requires inoculation by an infected leafhopper. If this is the mode of transmission, the rate of transmission would be low, and may explain the severity of infection due to random exposure in nature. The analyses of the infection pathway in leafhopper salivary glands (*Psammotettix alienus* Dahlbom), along with the capacity to induce 100% mortality in leafhopper cell cultures [*Homalodisca vitripennis* (Germar)] suggests Taastrup-like virus may have an early evolutionary association with leafhoppers. The rapid mortality observed in leafhopper cells when inoculated may also be why infected leafhoppers are rare, and difficult to collect in field sampling efforts.

## Acknowledgements

The authors are indebted to Prof. David B. Collinge for helpful advice during the preparation of the manuscript. Critical reviews by Dr. Xiomara Sinisterra-Hunter, AgTec, Plant Biotechnology, LLC and Dr. Catherine Katsar, PPQ, APHIS; Diann Achor, for electron microscopy work, University of Florida, IFAS, Lake Alfred, FL. Maria T. Gonzalez, Biological Science Technician, USDA, ARS, Fort Pierce, FL, for technical support.

